# Differential alteration of IL-8 in liver cancer stem cell enrichment in response to PI3K/Akt/mTOR inhibitors and sorafenib

**DOI:** 10.1101/306910

**Authors:** Deniz Cansen Kahraman, Tamer Kahraman, Rengul Cetin Atalay

## Abstract

Liver cancer stem cells (LCSCs) are derived from damaged and transformed Hepatic progenitor cells (HPCs) during precancerous cirrhosis stage. Ras/Raf/MAPK and PI3K/AKT/mTOR signaling pathways are significantly deregulated in liver cancer. The activation of PI3K/AKT/mTOR pathway in LCSC population is one of the reasons for acquired resistance to Sorafenib in advanced Hepatocellular carcinoma (HCC) patients. Therefore, identifying novel inhibitors targeting this pathway acting on LCSCs is highly essential. We therefore elucidated the bioactivities of small molecule kinase inhibitors on LCSCs acting through PI3K/Akt/mTOR pathway in comparison with DAPT (CSC inhibitor), DNA intercalators and Sorafenib. For this purpose, CD133+/EpCAM+ cells originated from HCC cells were analyzed by flow cytometry and effective inhibitors on LCSCs were further tested for their potential combinatorial effects. Treatment of cells with Sorafenib, and DNA intercalators resulted in enrichment of CD133+/EpCAM+ cells. Yet, mTOR inhibitor Rapamycin, and Notch pathway inhibitor DAPT significantly reduced CD133/EpCAM positivity. Combination studies revealed that sequential treatment strategy, which involves treatment of cells with Rapamycin prior to Sorafenib treatment, decreased the ratio of LCSCs as opposed to Sorafenib treatment alone or Sorafenib treatment prior to Rapamycin. The effect of the inhibitors were also demonstrated with LCSC sphere formation. Additionaly, a large panel of genes involved in cancer pathways were analyzed using Nanostring^®^ nCounter^®^ Technology to identify the differentially expressed genes in Rapamycin, Sorafenib or DAPT treated cells. Pathways involved in stemness (Wnt and Notch pathways) were differentially regulated between Rapamycin or DAPT treated cells and Sorafenib treated cells. Interleukin 8 (IL-8), FLNC, FLNA expressions were down-regulated upon treatment with DAPT or Rapamycin, yet up-regulated upon Sorafenib treatment. Following IL-8 inhibition CD133/EpCAM positivity of cells decreased significantly, indicating that IL-8 signaling is crucial for the conservation of stemness features of cancer cells.

**Conclusion:** PI3K/Akt/mTOR pathway inhibitors alter hepatic CSC composition and gene expression in favor or to the detriment of cancer stem cell survival. Blockade of IL-8 signaling provides a promising therapeutic approach for prevention of LCSC enrichment.

## Introductory Statement

Hepatocellular carcinoma (HCC) is the second common cause of cancer mortality and the fifth most common malignancy in human cancers (1). Chronic liver injury is caused by viral diseases, exposure to chemicals, and other environmental or autoimmune conditions which constitute the risk factors for HCC (2, 3). Liver cancer is known to originate from hepatic progenitor cells (HPC) during cirrhosis and has a very heterogeneous histological structure. In this process, HPCs are damaged and transformed to liver cancer stem cells (LCSCs) which are responsible for chemo-resistance, tumor relapse and metastasis (4). Sorafenib and Regorafenib are multikinase inhibitors, which are the only clinical treatment options for advanced HCC patients approved by Food and Drug Administration (FDA). However, in recent years it has been reported that HCC patients acquire resistence to Sorafenib. The underlying mechanisms of acquired Sorafenib resistance involve crosstalks of PI3K/Akt and JAK-STAT pathways, epithelial-mesenchymal transition (EMT), as well as enrichment of cancer stem cells (5, 6). Nevertheless the exact mechanisms accounting for sorafenib resistance remains unclear. Therefore, therapies targeting cancer stem cells in liver cancer will be indispensable for the treatment of HCC.

Markers validated for the identification of LCSCs include CD133, EpCAM, CD90, CD44, CD24, CD13, and OV6 as well as Hoechst dye efflux or aldehyde dehydrogenase (ALDH) activity (7). In our previous study, it was well defined that CD133 and EpCAM were convenient to detect LCSCs of epithelial like (well-differentiated) cells, whereas CD90 was efficient to detect LCSCs of mesenchymal like (poorly differentiated) cells (8, 9).

We report here that the small molecule inhibitor Rapamycin inhibits the enrichment of LCSCs and regulates pathways related with stemness. Additionally, gene expression analysis along with pathway scoring results have revealed several potential targets such as IL-8 against cancer stem cell enrichment, where blockade of IL-8 pathway was shown to prevent enrichment of LCSC population significantly.

## Experimental Procedures

### Cell Culture

Huh7, Hep3B and, Mahlavu cells were maintained in Dulbecco’s Modified Eagle’s Medium (DMEM) (Invitrogen GIBCO), supplemented with 10% fetal bovine serum (FBS) (Invitrogen GIBCO), and 0.1 mM nonessential amino acid, whereas SNU475 cells were maintained in RPMI (Invitrogen GIBCO), supplemented 10% fetal bovine serum (FBS) and 2mM L-glutamine. Both media contained 100 units/mL penicillin and 100 units/mL streptomycin. Cells were grown at 37 °C in a humidified incubator under 5% CO_2_.

### Flow cytometry

HCC cells were inoculated into 100mm^2^ culture dishes (100.000-200.000 cells). 24 hours later, cells were treated with the compounds (IC_50_ conc.) for 72 hours. Dead cells that no longer remained attached to the surface of the culture plates were discarded through vacuum aspiration and cells that remained attached were collected to be fixed with 4% paraformaldehyde for 30 minutes. Huh7 and Hep3B cells were stained for cancer stem cell markers using anti-CD133/1 (AC133)-Biotin (Miltenyi, 130-090-664), anti-biotin-PE (Miltenyi, 130-090-756), anti-EpCAM-FITC (Miltenyi, 130-080-301), whereas Mahlavu and SNU475 cells were stained using anti-CD90-FITC (Miltenyi, 130-095-403). For isotype controls, mouse IgG1 isotype control-FITC conjugate (Miltenyi, 130-092-213), mouse IgG1 isotype control antibody-biotin conjugate (Miltenyi, 130-093-018) were used. Staining of cells was done according to the manufacturer’s protocol. Cells were analyzed using BD Accuri C6 Flow Cytometer and Software (BD Biosciences) or Novocyte flow cytometer and NovoExpress software (Acea Biosciences). The same staining procedure was applied for the analysis of HCC cells to determine the CSC marker positivity. For the detection of stemness marker expressions permeabilization of cells with 90% ice-cold methanol was done for 10 min prior to blocking. Nanog (1:400; cat# 3580, Cell Signaling) and OCT4 (1:200; cat#2750, Cell Signaling) antibodies and anti-rabbit IgG-Alexa 647 (1:2000, cat# ab150083, Abcam) were used.

### Isolation of LCSCs

For separation of CD133+/EpCAM+ cells from Huh7 and Hep3B cells, FACSMelody Cell Sorter (BD) was used. Anti-CD133/1(AC133)-Biotin (cat.no.130-090-664) and anti-biotin-PE (cat. no.130-090-756) was used to label CD133+ cells. anti-EpCAM-FITC (cat. no.130-080-301) was used to label EpCAM+ cells. Purity of separated cells were assessed using flow cytometry.

### Sulforhodamine B (SRB) cytotoxicity assay

All cells were inoculated into 96-well plates (1000-2000 cells/well). After 24 hours, cells were treated with the inhibitors and DMSO control in concentrations 40μM to 2.5μM in serial dilutions. After 72h of treatment, cells were fixed by cold 10% (w/v) trichloroacetic acid (MERCK) for an hour. Then the cells were washed with ddH2O and left to dry. For staining of the proteins, 50 μl of 0.4% SRB dye (Sigma-Aldrich) was applied to each well to be incubated at RT for 10 min. To remove extra staining, wells were washed with 1% acetic acid for several times and left for air-drying. SRB dye was solubilized using 100 μl/well 10 mM Tris-Base solution and absorbance was measured at 515 nm using a plate reader (ELx800, BioTek). The experiment was performed in triplicates and the absorbance values were normalized to DMSO controls.

### Sphere formation assay

Sphere formation from cells was triggered using in DMEM/F12 serum-free medium (Gibco, Invitrogen) supplemented with epidermal growth factor (20 ng/ml; Thermo Fisher), basic fibroblast growth factor (10 ng/ml; Sigma), B27 supplement (1:50; Invitrogen), Heparin (2 μg/ml; Sigma), insulin (5 μg/ml; Sigma), hydrocortisone (0.5 μg/ml; Sigma) and 100 units/mL penicillin and streptomycin (Gibco, Invitrogen). The cells were cultured in ultra-low attachment 96-well plates for 6-12 days.

### Analysis of Gene Expression (Nanostring^®^, nCounter^®^ Technology)

Huh7 cells were seeded in 100 mm culture dishes (200,000 cell/dish) and treated with the corresponding IC_50_ concentrations of the inhibitors or with control (DMSO) for 72h (72h for each inhibitor included in sequential treatment).

For extraction of RNA, RNeasy Mini Kit (Qiagen) was used as instructed by the manufacturer. The total RNA content of samples was measured using NanoDrop One spectrophotometer (Thermo Fisher Scientific). 50 ng RNA per sample was used for processing the nCounter^®^ PanCancer Pathway Panel gene expression analysis (Nanostring^®^). This technology contains 730 different probes binding complementary to corresponding target mRNAs (hybridization at 65°C overnight). The fluorescence barcodes specific for each gene are attached to each probe, enabling digital counting of each gene. Sample preparation was done with nCounter^®^ PrepStation and microscopy scanning of the barcode signals were done with nCounter^®^ Digital Analyzer (Nanostring^®^) as instructed by the manufacturer. Quality control, normalization and differential expression analysis were done using Nanostring nSolver^™^ 3.0 Analysis software together with its Advanced Analysis plug-in.

### Cellular pathway analysis

A signal transduction score flow algorithm for cellular pathway analysis called “Pathway Scoring Application” (10) was used to utilize the expression data obtained from Nanostring nCounter technology and evaluate the activity of cell signaling pathways. For this, the log2 differential expression values were transformed into ranked values by first sorting these values in ascending order and by ranking each log2 value with reference to all log2 values present in the gene list. The entrez gene ids of each gene were inserted next to the gene names. The ranked list was saved in txt format which contained entrez gene ids, gene name and the ranked gene expression values. Then this list was uploaded to Cytoscape 3.5 software platform to integrate the pathway scoring application. “Pathways in cancer (human)” KEGG pathway was uploaded to analyze the expression data. The enrichment scores assigned by the algorithm describes the biological response of a specific process (in this case treatment with different inhibitors) in the pathway.

### Quantitative RT-PCR (qRT-PCR)

Total RNA was isolated from cells using RNeasy RNA-purification kit (Qiagen) and RevertAid First Strand cDNA synthesis kit (Thermo Scientific) was used for cDNA synthesis. The primer sequences specific for the stemness markers (Nanog and Sox2), IL-8, FLNA, Jag1 and the internal control gene GAPDH used for qRT-PCR are; *IL-8 forward primer*: 5’ TCCTGATTTCTGCAGCTCTGT 3’, *IL-8 reverse primer*: 5’ AAATTTGGGGTGGAAAGGTT 3’, *Nanog forward primer*: 5’ TACCTCAGCCTCCAGCAGATG 3’, *Nanog reverse primer:* 5’ TCTTCGGCCAGTTGTTTTTCTG 3’ and *GAPDH forward primer*: 5’ TATGACAACGAATTTGGCTAC 3’, *GAPDH reverse primer*: 5’ TCTCTCTTCCTCTTGTGCTCT 3’. Quantitative PCR was performed using Light Cycler^®^ 96 Real-Time PCR System (Roche) with LightCycler^®^ 480 SYBR Green I Master (Roche) under the following conditions: initial denaturation at 95 °C for 10 min, amplification for 45 cycles (95°C 10 sec followed by 60°C 10 sec and 72°C 10 sec), and melting at 95°C 10 sec followed by 65°C for 60 sec and 97°C for 1 sec. The experimental Ct (cycle threshold) was calibrated against GAPDH control product. All amplifications were performed in triplicate. The ΔΔCt method was used to determine the amount of target gene in inhibitor treated cells relative to that expressed by DMSO treated cells.

### Western Blot analysis

Bio-Rad protein electrophoresis (Mini-PROTEAN^®^ Tetra Cell Systems and TGX^™^ precast gels) and transfer system (Trans-Blot^®^ Turbo Transfer System) were used according to the manufacturer’s protocol for western blot analysis. For gel electrophoresis, 20-40 μg of protein was used per well. Proteins were transferred to a PVDF membrane. For immunoblotting, α-β-actin (Cell Signaling, 4967), α-SOX2 (Cell signaling, 2748S), α-NANOG (Cell signaling, 3580S) antibodies were used. Proteins were visualized using a C-Digit^®^ imaging system (Ll-COR).

### Statistical analysis

All data in this study were obtained from three independent experiments and S.D values were calculated accordingly. Statistical analysis was performed using a Student’s t-test (Prism, Graphpad or Microsoft Excel). MFI values of all flow cytometry analysis results were provided in **Suppl. fig. 5**.

## Results

### Isolated CD133+/EpCAM+ cells display cancer stem cell characteristics

CD133+/EpCAM+ and CD133-/EpCAM-populations from Huh7 and Hep3B cells were separated by cell sorter and analyzed by flow cytometry to determine the purity of the separated cells. The purity and sorting efficiency was ~95% for each population (Fig. 1A). To characterize the isolated cells in terms of their stemness features, sphere formation capacity and expression of well-known stemness factors (Nanog and Oct4) were compared between CD133+/EpCAM+ and CD33- /EpCAM- populations. It was found that, cells that were positive for both markers were able to form vivid and dense spheres, whereas cells that were negative for both markers, were unable to form such spheres (Fig.1B). In addition, the expression of Nanog and Oct4 were found to be higher in CD133+/EpCAM+ cells when compared to CD133-/EpCAM- cells (Fig. 1C). Altogether, these results indicated that CD133 and EpCAM markers were convenient for identifying cells that have cancer stem cell like features in epithelial-like HCC cell lines and were compatible with the literature (11, 12).

**Fig. 1.**
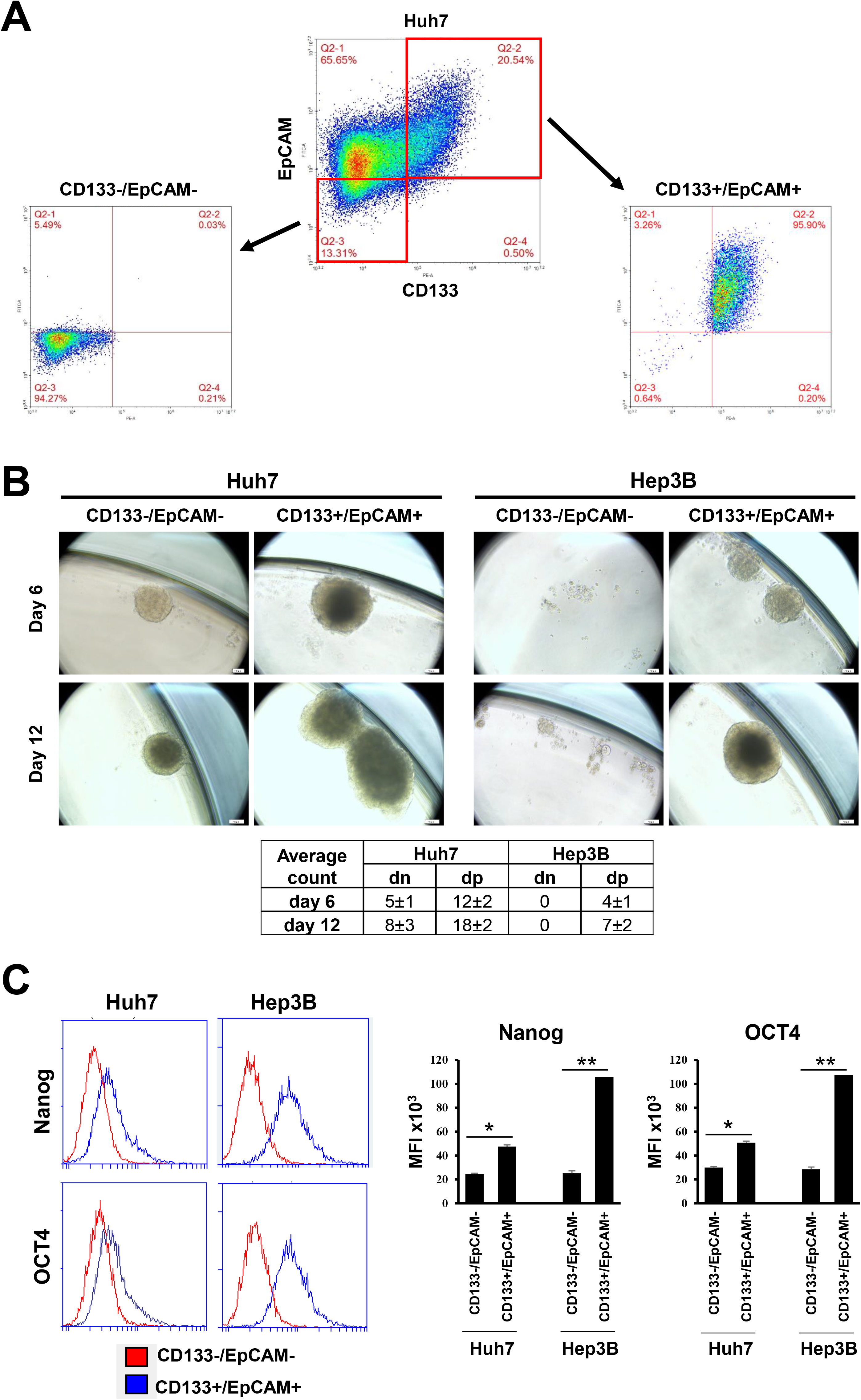
Characterization of Liver Cancer Stem Cells of HCC cells Huh7 and Hep3B. **(A)**. Flow cytometry analysis representing purity of CD133-/EpCAM- and CD133+/EpCAM+ populations of Huh7 cells after cell sorting. **B.** Sphere formation potential of CD133+/EpCAM+ population compared to CD133-/EpCAM- population in Huh7 and Hep3B cells on day 6 and day 12. Isolated populations of Huh7 and Hep3B cells were inoculated into low-attachment plates (200 cell/well) in liver cancer sphere formation media (40) and incubated until the spheres observed at 37 °C in a humidified incubator under 5% CO_2_. Microscopic images are representative of spheres formed by each population, and the sphere counts together with standard deviation values in double negative (dn) and double positive (dp) cells for CD133 and EpCAM markers **C.** Expression of stemness markers (Nanog and OCT4) assessed by fluorescence staining and flow cytometry analysis compared in CD133+/EpCAM+ (blue) and CD133-/EpCAM- populations (red) of Huh7 and Hep3B cells. x-axis represents the fluorescence intensity, while y-axis indicates cell count.

### Small molecule inhibitors have differential effects on LCSC population

A group of small molecule inhibitors which include several PI3K/Akt/mTOR pathway inhibitors, DNA intercalators, and Sorafenib were chosen to determine their effects on liver cancer stem cells. Initially, the inhibitory concentrations (IC_50_ values) of these inhibitors on HCC cells (Huh7, Hep3B, Mahlavu and SNU475) were evaluated by NCI-SRB assay. The IC_50_ values showed small differences between cell lines, therefore an average IC_50_ value was assigned to each inhibitor (Table 1). These concentrations were used throughout the following experiments explained in this study.

**Table 1:** Small molecule kinase inhibitors and their known target/s used in this study.

| **Inhibitor** | **Target/s** | **IC<sub>50</sub> (μM)** |
| --- | --- | --- |
| **Sorafenib** | B-Raf, VEGFR | 8.0 |
| **ZSTK474** | PI3K | 7.0 |
| **PI-103** | PI3K, mTOR | 4.0 |
| **PI3Ki-α** | PI3K-α | 0.1 |
| **PI3Ki-β** | PI3K-β | 40.0 |
| **AKTi-2** | Akt2 | 20.0 |
| **AKTi-1,2** | Akt1, Akt2 | 5.0 |
| **LY294002** | PI3K | 4.0 |
| **Wortmannin** | PI3K | 20.0 |
| **Rapamycin** | PI3K, mTOR | 0.1 |
| **BEZ-235** | PI3K, mTOR | 1.0 |
| **Camptothecin** | DNA intercalator | 0.1 |
| **Doxorubicin** | DNA intercalator | 0.1 |
| **DAPT** | γ-secretase-Notch | 5.0 |

*Average IC_50_ values of each inhibitor on liver cancer cells. Treatments with all inhibitors were done in triplicates. Average Standard deviations were less than 10% for all values.

** DNA intercalators, Camptothecin and Doxorubicin, and Notch inhibitor DAPT were used as controls in this study.

To observe the changes in the expression of stem cell markers after treatment with each inhibitor, cells were treated with the corresponding IC_50_ concentrations and incubated for 72 h. Dead cells were removed and the cells remaining alive were collected to be stained and analyzed by flow cytometry. Percentage of cells that were positive for both CD133 and EpCAM were compared between samples with reference to control group (DMSO). The results have shown that, in parallel with the literature, treatment of HCC cells with Sorafenib, and DNA intercalators resulted in enrichment of CD133+/EpCAM+ cells. PI3K/Akt pathway inhibitors led either to the enrichment of CD133+/EpCAM+ cells (ZSTK474, NVP-BEZ235) or did not have any effect (Akti-1,2, Akti-2, PI3Ki-α, PI3Ki-β) on LCSC populations. However, mTOR inhibitor-Rapamycin, PI3K inhibitor-LY294002 and Notch inhibitor-DAPT significantly reduced the CD133+/EpCAM+ cell population (Fig. 2). Yet, since LY294002 displayed its effect in lower significance, Rapamycin was selected to be included in combination experiments with Sorafenib.

**Fig. 2.**
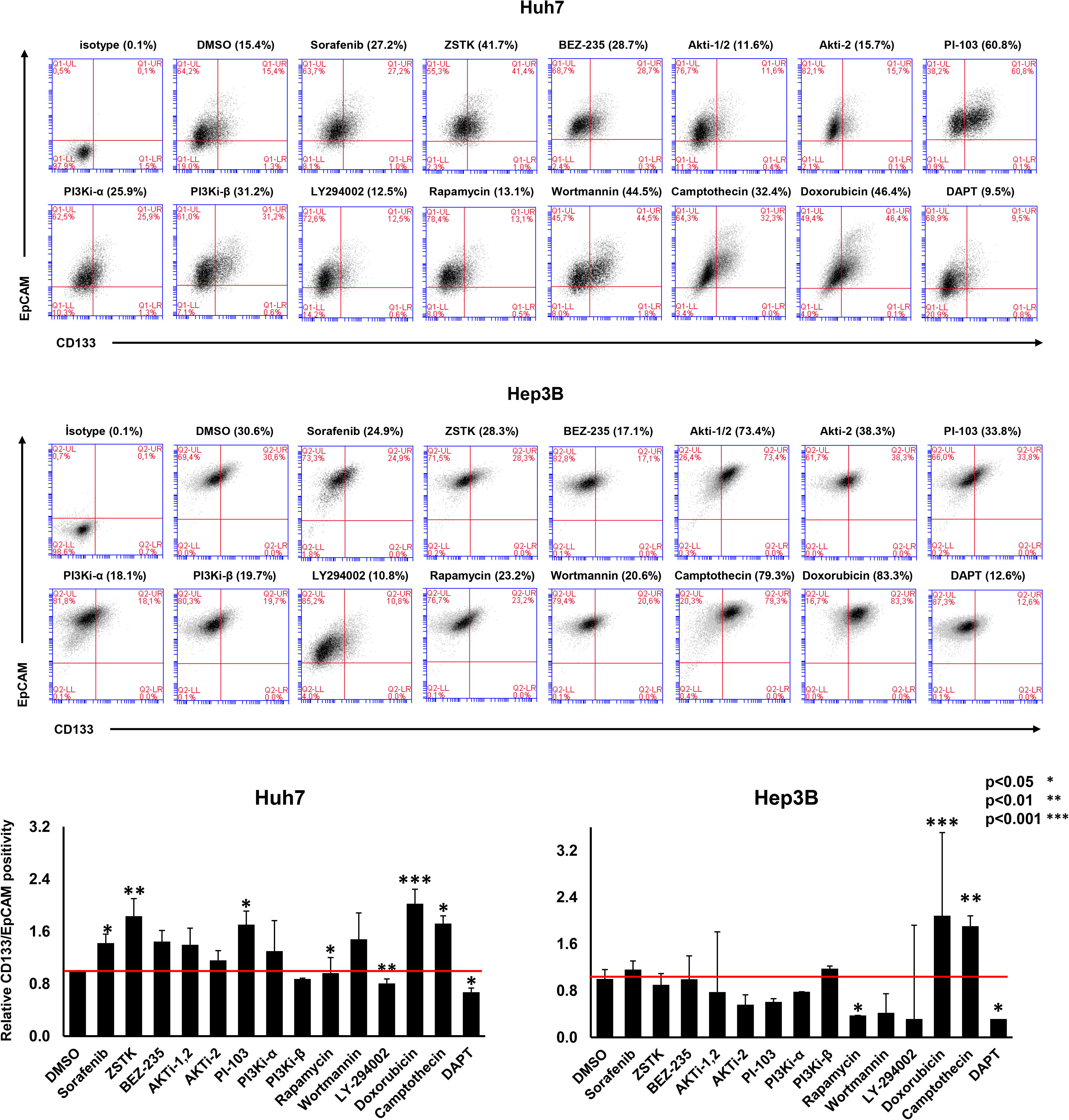
Effects of small molecule inhibitors on Liver Cancer Stem Cell population from Huh7 and Hep3B cells. Representative flow cytometry results indicating positivity of CD133/EpCAM population after 72 hours of treatment with IC_50_ concentrations of small molecule inhibitors (Table 1) vs. control vehicle (DMSO). Cells that survive each treatment were collected and stained fluorescently by anti-CD133-biotin, anti-biotin-PE and EpCAM-FITC antibodies. x-axis indicates CD133 positivity, and y-axis indicates EpCAM positivity. Lower-left quadrant, CD133-/EpCAM-; Upper-left quadrant, CD133-/EpCAM+; Lower-right quadrant, CD133+/EpCAM-; upper-right quadrant, CD133+/EpCAM+. Each treatment was compared to its corresponding DMSO control to define the changes in percentage of double positive population. DAPT was used as positive control for CSC inhibition. Bar graphs represent relative CD133/EpCAM positivity with respect to DMSO control group (red line: threshold). Inhibitors that show their effect significantly were indicated with * if p<0.05, ** if p<0.01, *** if p<0.001. Experiments were done in 4 biological replicates at different times.

Additionally, the changes in the cancer stem cell positivity of mesenchymal-like cells (Mahlavu and SNU475) were also determined when treated with Rapamycin, Sorafenib, Doxorubicin and DAPT. It was shown that Sorafenib and Doxorubicin significantly causes increase in CD90 positivity, on the contrary, Rapamycin and DAPT were able to decrease the same population (**Suppl. Fig. 1**). Altoghether, it was concluded that, Rapamycin was able to prevent enrichment of CSCs both in epithelial and mesenchymal like HCC cell lines similar to the effect of a well known cancer stem cell inhibitor DAPT.

### Rapamycin treatment before Sorafenib is effective against LCSC enrichment

Recent studies have introduced that mTOR plays critical roles in breast, pancreatic, lung and colorectal cancer stem cells through stemness related functions and that inhibition of mTOR significantly causes sensitization of cancer stem cells to therapy (13–16). It has already been shown by previous studies that Sorafenib synergizes with Rapamycin and prevents tumor progression in vivo (17) and that components of PTEN/AKT/mTOR pathway might be crucial in driving recurrence and effecting prognosis of HCC (18). However, the potential of combinatorial treatment using PI3K/AKT/mTOR pathway inhibitors and Sorafenib remain unclear. Therefore, two different sequential treatment strategies were followed; treatment of cells with 1. Sorafenib or Doxorubicin before Rapamycin, LY294002 or DAPT, 2. Treatment of cells with Rapamycin or DAPT before Sorafenib. Results have shown that the first strategy failed to inhibit enrichment of LCSCs and even enhanced this process (**Suppl. Fig. 2**). However, with the second strategy, it was possible to decrease the percentage of CD133+/EpCAM+ cells significantly in both Huh7 and Hep3B cells (Fig. 3) To further clarify the potential of this strategy, sphere formation assay was performed, which resulted in significantly smaller and low number of spheres in cells treated with Rapamycin prior to Sorafenib (Fig. 4). Thus, it was concluded that this strategy was promising in terms of inhibiting enrichment of LCSCs compared to Sorafenib treatment alone. Perhaps this effect was instigated by the regulation of stemness related events by the PI3K/AKT/mTOR pathway, which needs to be further investigated.

**Fig. 3.**
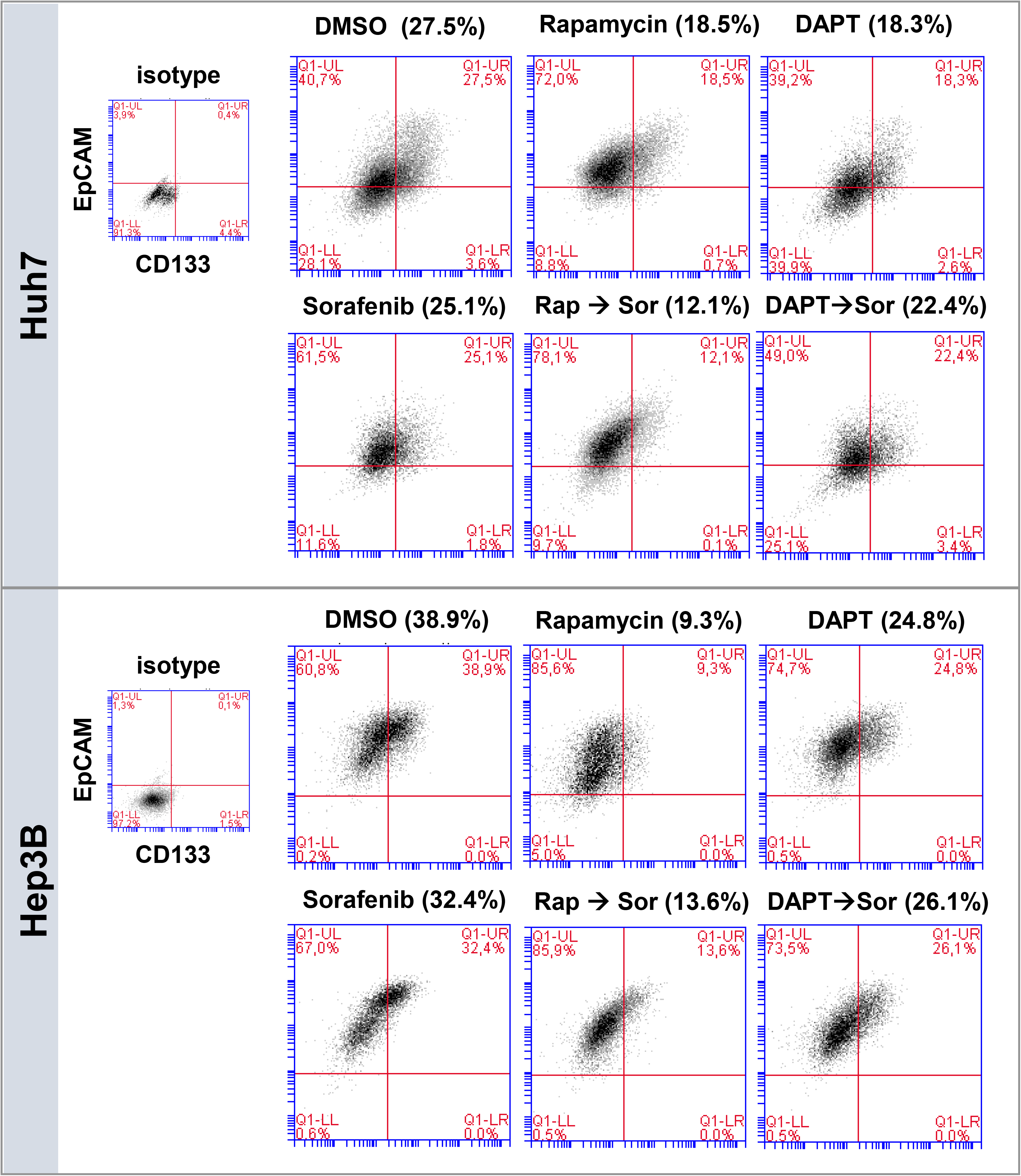
Positivity of CD133/EpCAM cells after sequential treatment strategy determined by flow cytometry analysis. Pre-treatment of Huh7 and Hep3B cells with IC_50_ concentrations of Rapamycin or DAPT for 72h followed by Sorafenib treatment (IC_50_ conc. for 72h Table 1). Cells that remain from each treatment group were collected and stained for CD133 and EpCAM using anti-CD133-biotin, anti-biotin-PE and EpCAM-FITC antibodies and analyzed with flow cytometry. Percent positivity of LCSCs in each group were compared between with respect to their DMSO controls. Rapamycin treatment before Sorafenib decreased number of cells positive for CD133 and EpCAM in both Huh7 and Hep3B cells compared to cells treated with Sorafenib only.

**Fig. 4.**
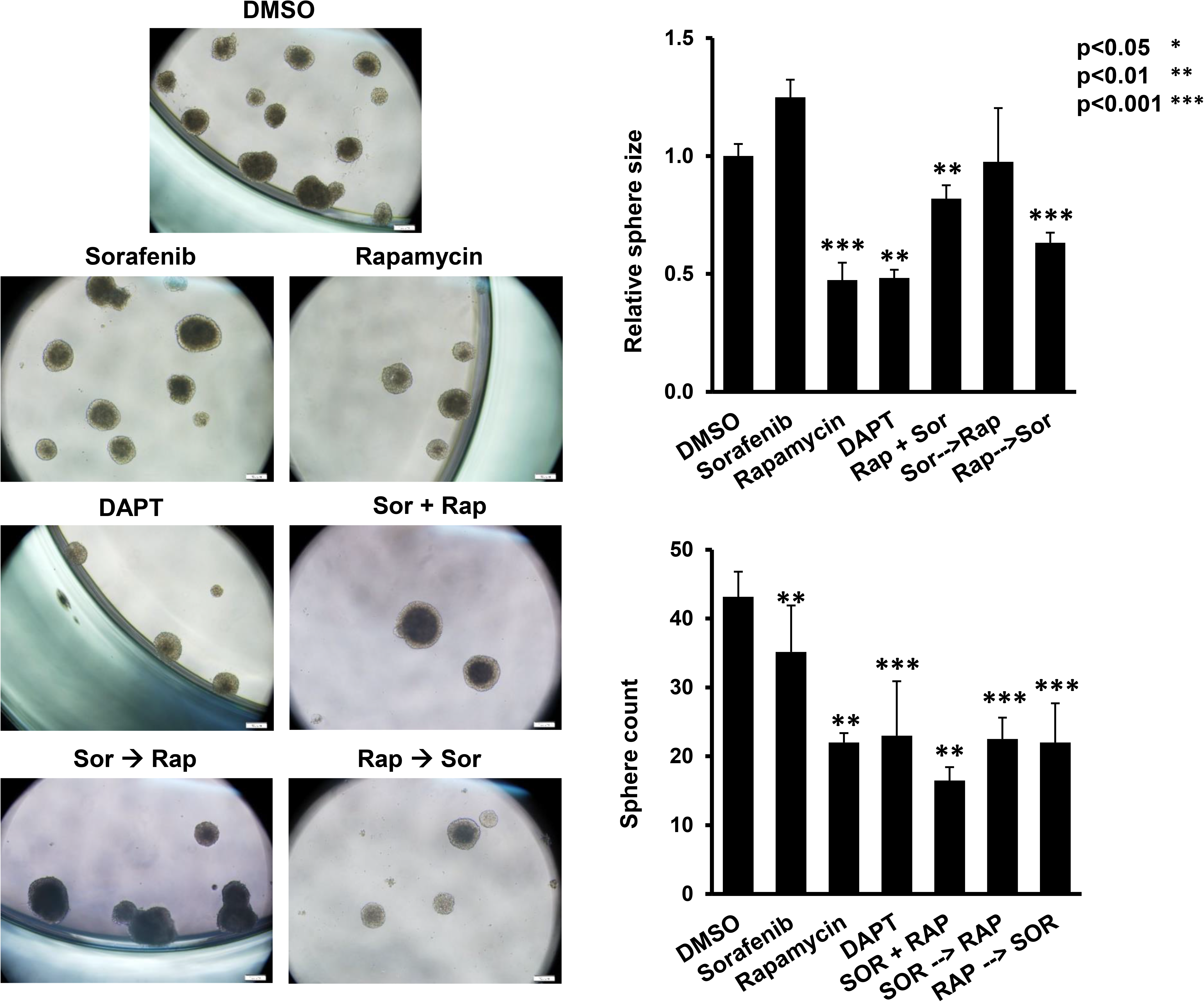
Sphere formation capacity of Huh7 cells. Microscopy images represent spheres formed after 6 days by Huh7 cells treated with either a single inhibitor (IC_50_ conc. for 72h) or in combination (corresponding IC_50_ conc. for each inhibitor were used for 72h at the same time (Sor+Rap) or sequentially (Rap 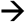 Sor or Sor 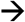 Rap)) and inoculated into ultra-low attachment plates in sphere formation media to be incubated in 37 °C in a humidified incubator under 5% CO_2_ until spheres were formed. Bar graphs represent sphere size and sphere count at day 6 for each treatment relative to DMSO treated group. Significance of results for each treatment group relative to DMSO were indicated as follows: * p<0.05, ** p<0.01, *** p<0.001.

### RNA expression analysis and pathway scoring results suggest potential targets against enrichment of LCSCs

To compare the effects of Sorafenib, Rapamycin and DAPT on expression of cancer pathway and stemness related genes, PanCancer pathways panel was analyzed with the help of Nanostring^®^ nCounter^®^ Technology. This platform offers high level of precision and sensitivity with its novel digital color-coded barcode technology as well as higher dynamic range of detection compared to microarrays and high-throughput sequencing technologies. (19, 20) Differential expression analysis have revealed that upon Rapamycin or DAPT treatment, stemness pathway genes (WNT and Notch) were down-regulated similarly (Fig. 5A), whereas Sorafenib treatment caused upregulation of JAG1 which encodes for Jagged-1, a Notch receptor ligand known to be responsible for enhancing cancer development and angiogenesis in cancer including liver cancer (21) (**Suppl. Fig. 3**). Similarly, when top 15 differentially expressed genes were clustered, expression of FLNA, FLNC genes were upregulated upon Sorafenib treatment. Filamin A has been indicated as a potential marker for the progression (metastasis) of HCC and other cancers (ovarian and prostate) in previous studies (22–24) (Fig. 5B). Once cells were treated with Rapamycin, DAPT or Rapamycin and then Sorafenib, the expression of these genes were found to be regulated inversely. Perhaps the most important finding was that, IL-8 expression was strongly upregulated (~5 fold, log2) upon Sorafenib treatment but on the contrary downregulated upon treatment with DAPT (~-3 fold, log2), or not changed when treated with Rapamycin (~0.18 fold, log2). Further confirmation of these results was done with qRT-PCR where gene expression levels of IL-8 were in parallel with the results obtained from the data obtained from the nCounter system (Fig. 5C). Nanog expression was also shown to be in parallel to expression of IL-8 in Huh7 cells and decreased upon Rapamycin treatment (Fig. 5C). In addition, there was a strong correlation between the ratio of LCSCs and expression of IL-8 in these cells. As the LCSC population is enriched, the expression of IL-8 gene increased likewise (Fig. 5D and **Suppl. Fig 4**). IL-8 gene, encoding for the cytokine IL-8, has been widely studied in many cancers including breast (25–28), lung (29),pancreatic (30), colorectal (31, 32), HCC (33), and has been shown to be associated with cancer stem cell-like properties. Therefore, targeting IL-8 signaling was found to be critical for the inhibition of CSC regulated events.

**Fig. 5.**
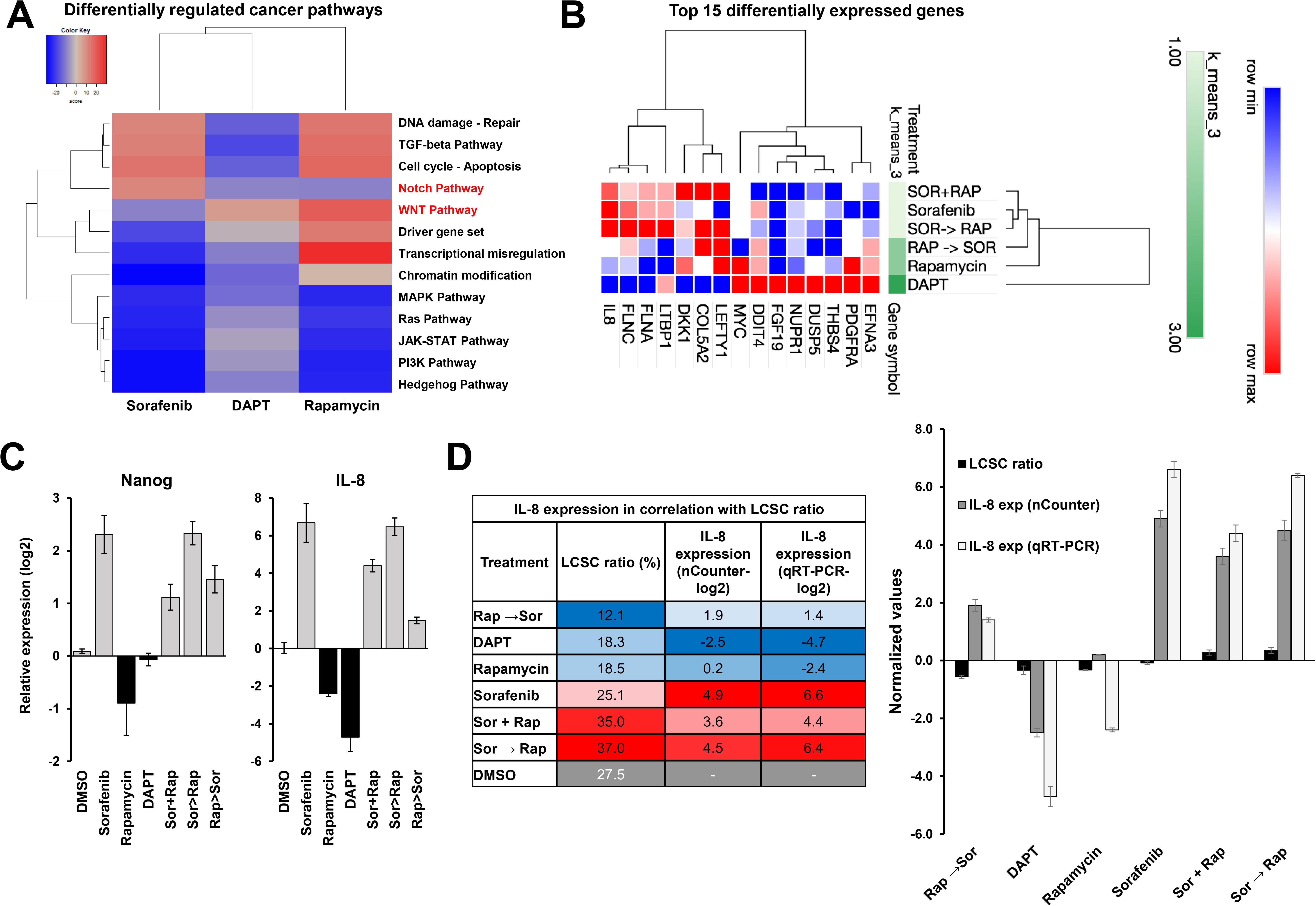
Gene expression analysis and pathway scoring results. **A**. 20 of the most significant differentially expressed genes in PanCancer pathways panel of Nanostring^®^ nCounter^®^ Technology with reference to DMSO control group. **B**. Heatmap representing the clustering of three different treatment groups (Sorafenib, DAPT and Rapamycin) with respect to differentially expressed genes involved in different cancer pathways **C**. Bar graph demonstrating relative expression of IL-8 obtained by qRT-PCR. **D**. Score flow algorithm results indicating relationship of genes involved in IL-8 signaling. The scores for each gene are represented as follows; high (red), intermediate (yellow) and low (green).

The differential expression data obtained was further used to execute a score flow algorithm (“Pathway scoring application”) (10) (Fig. 6A) in order to evaluate the activity of cell signaling pathways. In fact, parallel with the differential gene expression analysis, IL-8 gene gained significantly enriched scores in Sorafenib treated cells compared to DAPT or Rapamycin treated cells, further emphasizing the potential of IL-8 as a target in HCC (Fig. 6B).

**Fig. 6.**
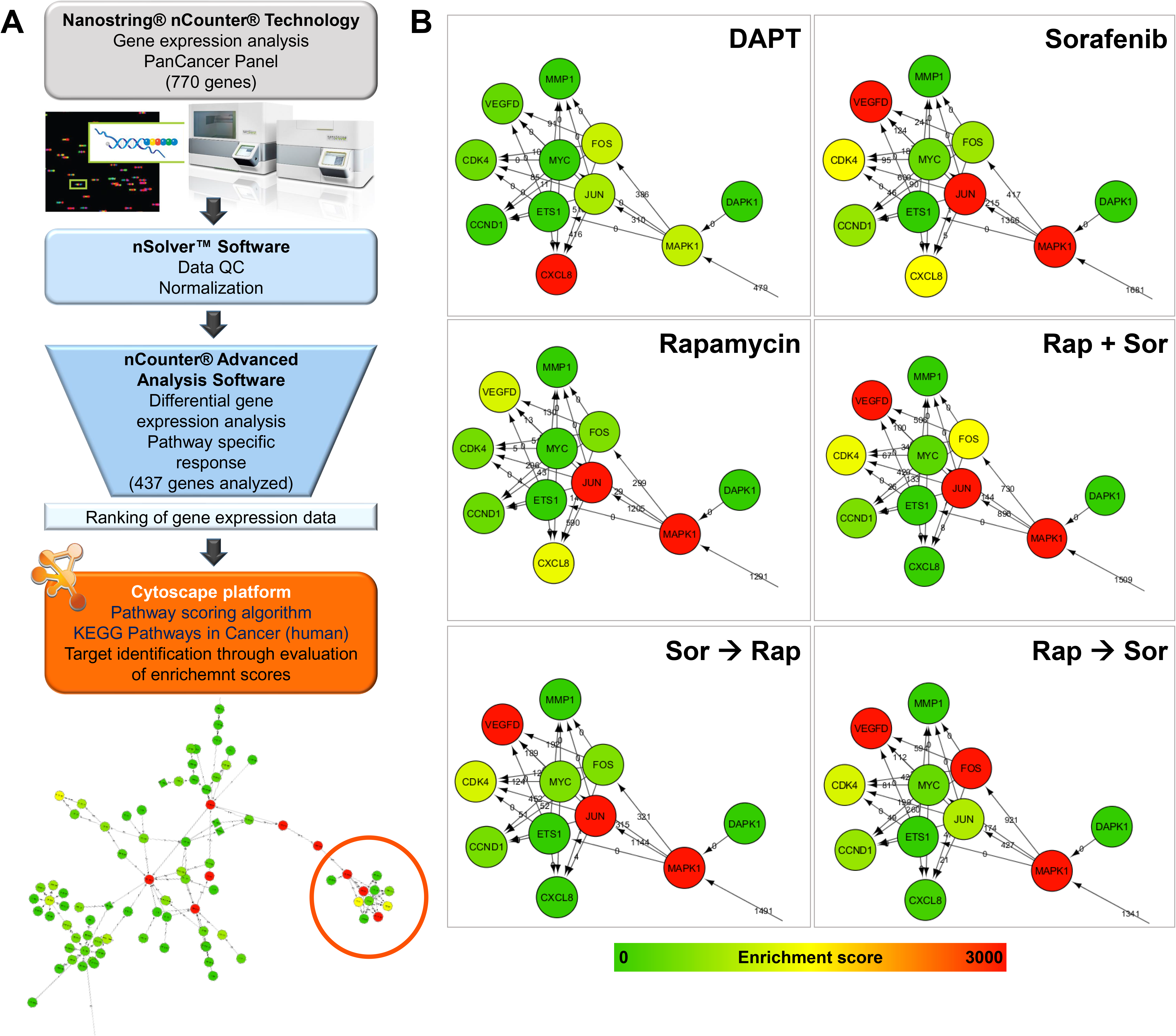
Pathway scoring results using differential gene expression data obtained from Nanostring nCounter platform. **A**. Flow chart of steps followed to adapt the differential gene expression data into our pathway scoring algorithm using Cytoscape platform and KEGG Pathways in Cancer (Homo sapiens) pathway. Initially, expression analysis of 770 genes involved in PanCancer pathway panel were done using the Nanostring nCounter technology and the analysis softwares available. After quality control and normalization of the expression data, advanced analysis software was used to obtain differentially expressed (DE) genes in each group relative to DMSO. Among 770 genes, 437 were able to be statistically computable. Then, differential expression results of each treatment group were converted into ranked lists which include gene name, entrez gene id and ranked DE value of each gene (as explained in the methods part) and loaded into Cytoscape 3.5 platform where “pathway scoring algorithm” plug-in was run integrated with “Pathways in cancer (human)” KEGG pathway. The enrichment scores obtained were represented with a color scale where scores increase as the color shifts from green to red. **B**. Pathway scoring results of CXCL8 (IL-8) and the surrounding effectors are shown for each treatment group.

### Interleukin 8 inhibition sensitizes HCC cells to Sorafenib treatment

Former results have lead us to concentrate on the possibility that HCC cells could be sensitized to Sorafenib treatment through blockade of IL-8 signaling. For this purpose, Reparixin, an IL-8 inhibitor, was used to inhibit IL-8 signaling in Huh7 and Hep3B cells. The IC_50_ value obtained by SRB assay was 50 μM and treatment of cells with Reparixin alone and in combination with Sorafenib, resulted in significant reduction in positivity of LCSC markers (Fig. 7A), as well as their capacity to form spheres (Fig. 7B, 7A and 7D). In addition, western blot analysis of these spheres has shown that both Nanog and SOX2 protein levels were lower in spheres formed by Huh7 cells treated with DAPT, Reparixin, whereas spheres formed by Sorafenib treated cells had higher levels of both stemness markers. However, the combination of Sorafenib and Reparixin resulted in lower levels of SOX2 and Nanog proteins (Fig. 7E). Altogether these results supported the idea that IL-8 signaling is associated with LCSC enrichment and that targeting IL-8 signaling is a promising in terms of coping with HCC pathogenesis.

**Fig. 7.**
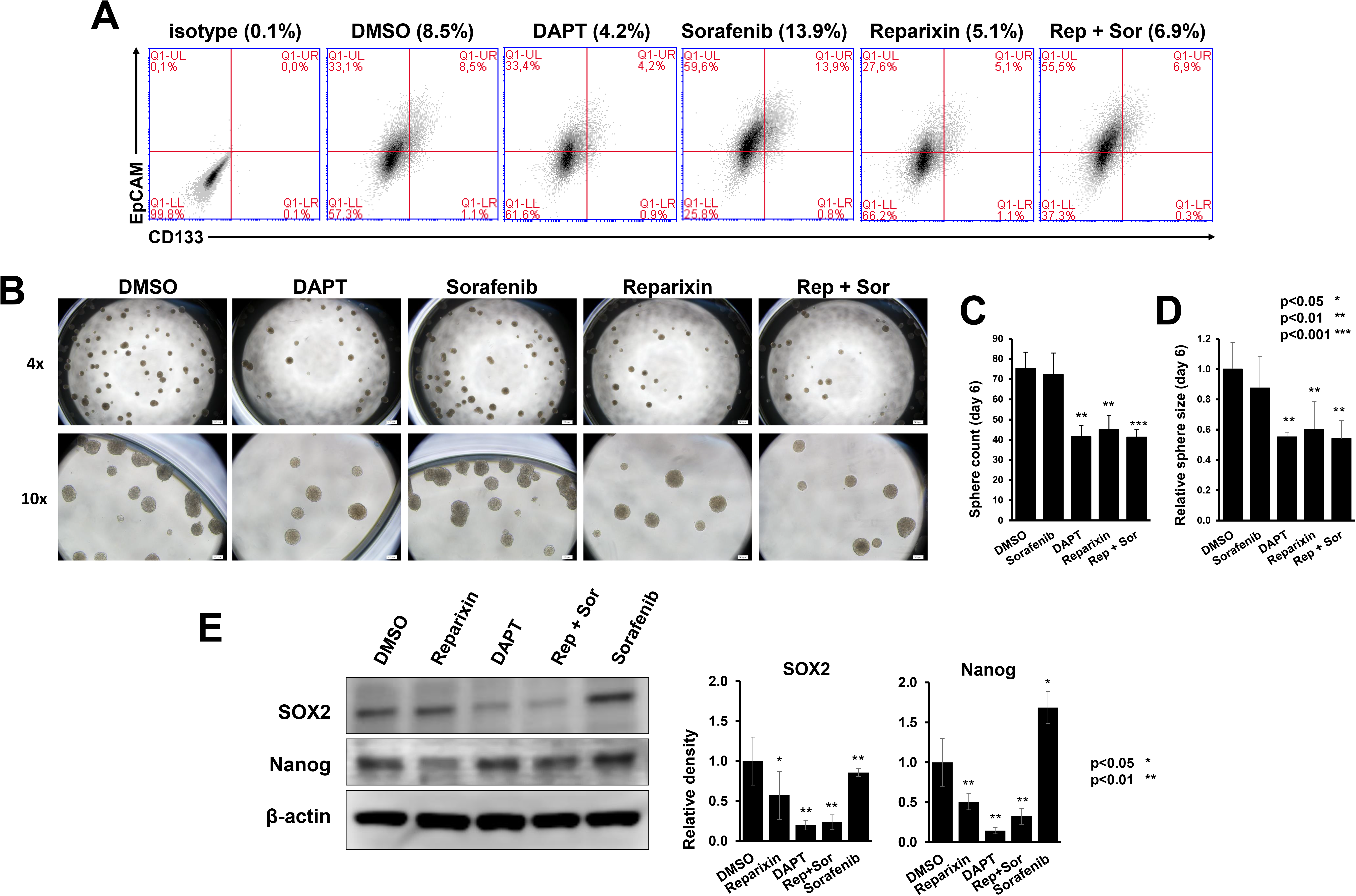
Effects of IL-8 inhibition by Reparixin on LCSC enrichment. **A**. Flow cytometry analysis of Huh7 cells treated with IC_50_ concentrations of indicated inhibitors for 72h. Table summarizing the % LCSC positivity for each treatment group **B**. Representative 4x and 10x microscopic images of spheres formed by Huh7 cells after 72h of treatment with IC_50_ conc. of indicated inhibitors. **C**. Bar graph comparing the number of spheres for each treatment group and **D**. bar graph representing sphere size relative to DMSO control. Significance of differences with respect to DMSO treated group were indicated as follows: * p<0.05, ** p<0.01, *** p<0.001. **E**. Western Blot analysis of whole lysate extracted from spheres of each treatment group. Representative images and bar graphs show protein levels of SOX2 and NANOG. β-actin was used as loading control and relative intensity values for both proteins were obtained using LI-COR Image studio software.

## Discussion

In this study characterization of CD133+/EpCAM+ population of Huh7 and Hep3B cells were done to demonstrate their cancer stem cell like features and to verify the efficacy of CD133 and EpCAM molecules as LCSC markers. Both sphere formation assay results and expression analysis of Nanog and OCT4 have once again supported the previous studies performed by other groups and showed that CD133 and EpCAM together are convenient to identify and isolate cancer stem cell like cells of HCC (Fig. 1). Supportive to our results, literature has also defined the association between PI3K/Akt/mTOR pathway and LCSC markers CD133 and EpCAM through analysis of tumor and non-tumor tissues of from HCC patients and found that p-AKT and p-mTOR expression was positively correlated with CD133, EpCAM and CD90 expression and together these markers were correlated with poor differentiation and advanced tumor stage (18). Thus, expression of PTEN in combination with CD133 or EpCAM was suggested to serve as a screening tool to monitor recurrence and predict prognosis in HCC.

Although Sorafenib and other small molecule kinase inhibitors targeting PI3K/AKT or Ras/Raf pathways are widely studied for their anti-cancer activity, it has become a great concern that their efficacy is unstable in HCC patients due to high-level heterogeneity. It is suggested by many studies that, since Sorafenib inhibits Raf/(MAPK) signaling but activates PI3K/Akt pathway (34), combinations targeting Akt/mTOR or Ras/MAK pathways could be promising in ensure better clinical outcomes. In fact, it has already been shown that PKI-587, an inhibitor of PI3K/AKT/mTOR pathway together with Sorafenib is advantageous in terms of inhibition of HCC (35). However, a recently completed phase II clinical trial (36) which compares everolimus (targeting PI3K/AKT/mTOR pathway) combined with Sorafenib, and a phase III trial of Sorafenib and erlotinib (an EGFR inhibitor) (37) showed no improved survival compared to Sorefenib alone (38). Given the fact that not all PI3K/AKT/mTOR pathway inhibitors exhibit promising effect on HCC, a group of novel/known small molecule inhibitors including PI3K/AKT pathway inhibitors, Sorafenib and clinically used chemotherapeutics (DNA intercalators) were investigated for their efficacy against LCSCs of HCC cell lines. As expected, and in parallel with the literature, each inhibitor was found to have differential effects on CD133+/EpCAM+ population where; Sorafenib and DNA intercalators such as Doxorubicin and Camptothecin and some PI3K/Akt pathway inhibitors (ZSTK and NVP-BEZ235) enriched the ratio of CSCs; whereas some inhibitors decreased the CSC population significantly (Rapamycin, and LY294002) (Fig. 2). Therefore, we claim that PI3K/Akt/mTOR pathway inhibitors may alter hepatic cancer stem cell composition and gene expression in favor or to the detriment of cancer stem cell survival, meaning that the composition and the timing of the combined treatment with Sorafenib can be in favor of the patient survival or a risk for tumor relapse. For this reason inhibitors that were shown to be potent against LCSCs were further tested in combination with Sorafenib to propose a treatment strategy that would get in the way of LCSC enrichment or sensitize them to become defenseless against Sorafenib treatment. Our findings showed that sequential treatment which involves treatment of cells with Rapamycin prior to Sorafenib was effective against enrichment of CSCs compared to Sorafenib treatment alone or in combination (Fig. 3). It was interesting that when these inhibitors switch orders, completely different results were reached, where CSCs were likely to enrich even more compared to Sorafenib treatment alone (**Suppl. Fig. 2**). Perhaps pre-treatment of cells with Rapamycin caused cellular events that regulate the pathways related with stemness, thus helped sensitization of CSCs to the following treatment. In fact, literature has supportive information about the potential of mTOR pathway inhibitors to block stemness related pathways. A recent study introduced a novel mTOR inhibitor named Anthracimycin, which was shown to be capable of suppressing proliferation of CSCs and non-CSCs equally well in Huh7 cells (39).

In order to understand the alterations in expressions of cancer pathway related genes and to identify novel molecular targets that could have a significant role in survival mechanisms of LCSCs, a panel consisting total of 700 genes was studied and analyzed using Nanostring^®^ nCounter^®^ technology (Fig. 5). The differential expression data obtained was further used to execute a score flow algorithm (“Pathway scoring application”) to evaluate the activity of various signaling pathways (Fig. 6). As a positive control for the inhibition of cancer stem cells, upon DAPT treatment, genes related with Notch pathway were down-regulated as expected. Differential expression analysis revealed that expression of Ras, MAPK pathway related genes (PDGFRA, EFNA1, EFNA3, VEGFA, RASA4) were down-regulated, whereas genes belonging to Wnt and Notch pathway (DKK1 and JAG1) were upregulated upon treatment with Sorafenib. Regulation of IL-8 draw our attention since IL-8 signaling in cancer has been widely studied in many cancers, especially in breast cancer and its knockout resulted in poor survival of CSCs. In fact, it has been reported previously that IL-8 levels increase during progression of HCC and plays role in development of distant metastasis. Another study has described that IL-8 plays an important role in driving HCC tumor growth, self-renewal and angiogenesis in CD133+ LCSCs (33). Interestingly, treatment of cells with Rapamycin prior to Sorafenib, prevented upregulation of IL-8 unlike other combination strategies, which further suggested that Rapamycin could be inducing cellular events that do not allow IL-8 expression to elevate, yet triggers a mechanism which inhibits survival of CSCs. In fact, inhibition of IL-8 through treatment with Reparixin was informative, since it resulted in reduction of CD133+/EpCAM+ population, their sphere formation capacity and the levels of stemness related proteins (Fig. 7). To fully understand the exact mechanisms behind the regulation of cellular pathways involved in LCSC survival and resistance to Sorafenib, further understanding of the pathways induced by mTOR and IL-8 signaling in LCSCs is necessary. We believe that this study has provided a better understanding of alterations in distribution of stem/non-stem cells in HCC cells upon treatment with different PI3K/Akt/mTOR pathway inhibitors and possible mechanisms and targets involved in regulation of LCSC survival.

## Acknowledgements

This work was supported by TUBITAK research grant #113S540 and Turkish Ministry of Development project #2016K121540.

## References

1. Stewart BW, Wild C, International Agency for Research on Cancer, World Health Organization. World cancer report 2014: xiv, 630 pages.

2. Baffy G, Brunt EM, Caldwell SH. Hepatocellular carcinoma in non-alcoholic fatty liver disease: An emerging menace. Journal of Hepatology 2012;56:1384–1391.

3. Sun BC, Karin M. Obesity, inflammation, and liver cancer. Journal of Hepatology 2012;56:704–713.

4. Yamashita T, Wang XW. Cancer stem cells in the development of liver cancer. J Clin Invest 2013;123:1911–1918.

5. Palmer DH. Sorafenib in Advanced Hepatocellular Carcinoma. New England Journal of Medicine 2008;359:2498–2498.

6. Zhai B, Sun XY. Mechanisms of resistance to sorafenib and the corresponding strategies in hepatocellular carcinoma. World J Hepatol 2013;5:345–352.

7. Sun JH, Luo Q, Liu LL, Song GB. Liver cancer stem cell markers: Progression and therapeutic implications. World J Gastroenterol 2016;22:3547–3557.

8. Kahraman DC, Hanquet G, Jeanmart L, Lanners S, Sramel P, Bohac A, Cetin-Atalay R. Quinoides and VEGFR2 TKIs influence the fate of hepatocellular carcinoma and its cancer stem cells. Medchemcomm 2017;8:81–87.

9. Yamashita T, Kaneko S. Orchestration of hepatocellular carcinoma development by diverse liver cancer stem cells. J Gastroenterol 2014;49:1105–1110.

10. Isik Z, Ersahin T, Atalay V, Aykanat C, Cetin-Atalay R. A signal transduction score flow algorithm for cyclic cellular pathway analysis, which combines transcriptome and ChIP-seq data. Molecular Biosystems 2012;8:3224–3231.

11. Wilson GS, Hu ZN, Duan W, Tian AP, Wang XM, McLeod D, Lam V, et al. Efficacy of Using Cancer Stem Cell Markers in Isolating and Characterizing Liver Cancer Stem Cells. Stem Cells and Development 2013;22:2655–2664.

12. Chan AWH, Tong JHM, Chan SL, Lai PBS, To KF. Expression of stemness markers (CD133 and EpCAM) in prognostication of hepatocellular carcinoma. Histopathology 2014;64:935–950.

13. Matsubara S, Ding Q, Miyazaki Y, Kuwahata T, Tsukasa K, Takao S. mTOR plays critical roles in pancreatic cancer stem cells through specific and stemness-related functions. Scientific Reports 2013;3.

14. Lai YH, Yu XP, Lin XH, He SY. Inhibition of mTOR sensitizes breast cancer stem cells to radiation-induced repression of self-renewal through the regulation of MnSOD and Akt. International Journal of Molecular Medicine 2016;37:369–377.

15. Carrasco-Garcia E, Lopez L, Aldaz P, Arevalo S, Aldaregia J, Egana L, Bujanda L, et al. SOX9-regulated cell plasticity in colorectal metastasis is attenuated by rapamycin. Scientific Reports 2016;6.

16. Xie LX, Sun FF, He BF, Zhan XF, Song J, Chen SS, Yu SC, et al. Rapamycin inhibited the function of lung CSCs via SOX2. Tumor Biology 2016;37:4929–4937.

17. Newell P, Toffanin S, Villanueva A, Chiang DY, Minguez B, Cabellos L, Savic R, et al. Ras pathway activation in hepatocellular carcinoma and anti-tumoral effect of combined sorafenib and rapamycin in vivo. Journal of Hepatology 2009;51:725–733.

18. Su RJ, Nan HC, Guo H, Ruan ZP, Jiang LL, Song YY, Nan KJ. Associations of components of PTEN/AKT/mTOR pathway with cancer stem cell markers and prognostic value of these biomarkers in hepatocellular carcinoma. Hepatology Research 2016;46:1380–1391.

19. Tsang HF, Xue VW, Koh SP, Chiu YM, Ng LPW, Wong SCC. NanoString, a novel digital color-coded barcode technology: current and future applications in molecular diagnostics. Expert Review of Molecular Diagnostics 2017;17:95–103.

20. Geiss GK, Bumgarner RE, Birditt B, Dahl T, Dowidar N, Dunaway DL, Fell HP, et al. Direct multiplexed measurement of gene expression with color-coded probe pairs (vol 26, pg 317, 2008). Nature Biotechnology 2008;26:709–709.

21. Capaccione KM, Pine SR. The Notch signaling pathway as a mediator of tumor survival. Carcinogenesis 2013;34:1420–1430.

22. Ai JZ, Huang HZ, Lv XY, Tang ZW, Chen MZ, Chen TL, Duan WW, et al. FLNA and PGK1 are Two Potential Markers for Progression in Hepatocellular Carcinoma. Cellular Physiology and Biochemistry 2011;27:207–216.

23. Bedolla RG, Wang Y, Asuncion A, Chamie K, Siddiqui S, Mudryj MM, Prihoda TJ, et al. Nuclear versus Cytoplasmic Localization of Filamin A in Prostate Cancer: Immunohistochemical Correlation with Metastases. Clinical Cancer Research 2009;15:788–796.

24. Bourguignon LYW, Gilad E, Peyrollier K. Heregulin-mediated ErbB2-ERK signaling activates hyaluronan synthases leading to CD44-dependent ovarian tumor cell growth and migration. Journal of Biological Chemistry 2007;282:19426–19441.

25. Singh JK, Simoes BM, Howell SJ, Farnie G, Clarke RB. Recent advances reveal IL-8 signaling as a potential key to targeting breast cancer stem cells. Breast Cancer Research 2013;15.

26. Chin AR, Wang SE. Cytokines driving breast cancer stemness. Molecular and Cellular Endocrinology 2014;382:598–602.

27. Singh JK, Simoes BM, Clarke RB, Bundred NJ. Targeting IL-8 signalling to inhibit breast cancer stem cell activity. Expert Opinion on Therapeutic Targets 2013;17:1235–1241.

28. Ginestier C, Liu SL, Diebel ME, Korkaya H, Luo M, Brown M, Wicinski J, et al. CXCR1 blockade selectively targets human breast cancer stem cells in vitro and in xenografts. Journal of Clinical Investigation 2010;120:485–497.

29. Liu YN, Chang TH, Tsai MF, Wu SG, Tsai TH, Chen HY, Yu SL, et al. IL-8 confers resistance to EGFR inhibitors by inducing stem cell properties in lung cancer. Oncotarget 2015;6:10415–10431.

30. Chen LY, Fan J, Chen H, Meng ZQ, Chen Z, Wang P, Liu LM. The IL-8/CXCR1 axis is associated with cancer stem cell-like properties and correlates with clinical prognosis in human pancreatic cancer cases. Scientific Reports 2014;4.

31. Hwang WL, Yang MH, Tsai ML, Lan HY, Su SH, Chang SC, Teng HW, et al. SNAIL Regulates Interleukin-8 Expression, Stem Cell-Like Activity, and Tumorigenicity of Human Colorectal Carcinoma Cells. Gastroenterology 2011;141:279–U382.

32. Wang JC, Wang YN, Wang SC, Cai JY, Shi JQ, Sui X, Cao Y, et al. Bone marrow-derived mesenchymal stem cell-secreted IL-8 promotes the angiogenesis and growth of colorectal cancer. Oncotarget 2015;6:42825–42837.

33. Tang KH, Ma S, Lee TK, Chan YP, Kwan PS, Tong CM, Ng IO, et al. CD133+liver tumor-initiating cells promote tumor angiogenesis, growth, and self-renewal through neurotensin/interleukin-8/CXCL1 signaling. Hepatology 2012;55:807–820.

34. Wang CM, Cigliano A, Delogu S, Armbruster J, Dombrowski F, Evert M, Chen X, et al. Functional crosstalk between AKT/mTOR and Ras/MAPK pathways in hepatocarcinogenesis Implications for the treatment of human liver cancer. Cell Cycle 2013;12:1999–2010.

35. Gedaly R, Angulo P, Hundley J, Daily MF, Chen CG, Evers BM. PKI-587 and Sorafenib Targeting PI3K/AKT/mTOR and Ras/Raf/MAPK Pathways Synergistically Inhibit HCC Cell Proliferation. Journal of Surgical Research 2012;176:542–548.

36. Koeberle D, Dufour JF, Demeter G, Li Q, Ribi K, Samaras P, Saletti P, et al. Sorafenib with or without everolimus in patients with advanced hepatocellular carcinoma (HCC): a randomized multicenter, multinational phase II trial (SAKK 77/08 and SASL 29). Annals of Oncology 2016;27:856–861.

37. Zhu AX, Rosmorduc O, Evans TR, Ross PJ, Santoro A, Carrilho FJ, Bruix J, et al. SEARCH: a phase III, randomized, double-blind, placebo-controlled trial of sorafenib plus erlotinib in patients with advanced hepatocellular carcinoma. Journal of Clinical Oncology 2015;33:559–566.

38. Chen J, Jin RN, Zhao J, Liu JH, Ying HN, Yan H, Zhou SJ, et al. Potential molecular, cellular and microenvironmental mechanism of sorafenib resistance in hepatocellular carcinoma. Cancer Letters 2015;367:1–11.

39. Hayashi T, Yamashita T, Okada H, Oishi N, Sunagozaka H, Nio K, Hara Y, et al. A Novel mTOR Inhibitor; Anthracimycin for the Treatment of Human Hepatocellular Carcinoma. Anticancer Res 2017;37:3397–3403.

40. Cao L, Zhou Y, Zhai B, Liao J, Xu W, Zhang R, Li J, et al. Sphere-forming cell subpopulations with cancer stem cell properties in human hepatoma cell lines. BMC Gastroenterol 2011;11:71.

